# Wing tags severely impair movement in African Cape Vultures

**DOI:** 10.1101/2020.08.28.271700

**Authors:** Teja Curk, Martina Scacco, Kamran Safi, Martin Wikelski, Wolfgang Fiedler, Ryno Kemp, Kerri Wolter

## Abstract

**Background:** The use of tracking technologies is key for the study of animal movement and pivotal to ecological and conservation research. However, the potential effects of devices attached to animals are sometimes neglected. The impact of tagging not only rises welfare concerns, but can also bias the data collected, causing misinterpretation of the observed behaviour which invalidates the comparability of information across individuals and populations. Patagial (wing) tags have been extensively used as a marking method for visual resightings in endangered vulture species, but their effect on the aerodynamics of the birds and their flight behaviour is yet to be investigated. Using GPS backpack mounted devices, we compared the flight performance of 27 captive and wild Cape Vultures (*Gyps coprotheres*), marked with either patagial tags or coloured leg bands.

**Results:** Individuals equipped with patagial tags were less likely to fly, travelled shorter distances and flew slower compared to individuals equipped with leg bands. These effects were also observed in one individual that recovered its flight performance after replacing its patagial tag by a leg band.

**Conclusions:** Although we did not measure the effects of patagial tags on body condition or survival, our study strongly suggests that they have severe adverse effects on vultures’ flight behaviour and emphasises the importance of investigating the effects that tagging methods can have on the behaviour and conservation of the study species, as well as on the quality of the scientific results.

## Background

When measuring a phenomenon, the mere process of recording it influences our comprehension of the phenomenon itself [1]. The rapid development of tracking technologies and miniaturised sensors (bio-logging) allows us to finally answer long standing hypotheses about animal behaviour while providing a fertile environment to tackle novel research questions [2]. GPS loggers and geolocators record animal movements, accelerometers, cameras and proximity sensors can inform us about detailed behaviours and interactions among individuals and internal sensors such as heart rate loggers can provide information about the animals’ physiology. Finally, additional sensors can report on the environmental conditions experienced by the animals, such as temperature, air or water pressure and water depth.

The detail and spatial coverage of these data and the fact that they can be collected remotely from wild free-ranging animals are unprecedented, and impossible to achieve using direct observation [3]. Therefore the use of bio-logging and especially tracking technology is fundamental for the study of animal movement and pivotal to ecological and conservation research. However, the potential bias caused by the process of data collection has been often underrated.

Researchers are aware that tagging devices might affect the behaviour and survival of animals and more and more studies are suggesting solutions to minimise their effects, despite these being difficult to estimate and account for when interpreting results. Several studies have shown that the behaviour of birds can be affected by the attached devices [4–6]. A study on migrating birds, highlighted how return rates of individuals carrying geolocators were significantly lower than in the control group [7]. Devices attached to birds were also shown to negatively influence reproduction, nesting, parental care and survival, to reduce flight speeds [8, 9], and to increase energy expenditure [4, 6]. Some studies demonstrated a positive effect of devices on foraging trip duration [6] whereas [10] reported shorter foraging trips for birds equipped with devices compared to the control group.

These behavioural changes can not only have a negative impact on the reproduction and survival of animals [6], but also bias our understanding of animal behaviour [4]. In fact, effects of tagging methods pose, beyond being a concern for animal welfare, a major problem because of their systematic bias on the data. So far, there has been little or no research on how tagging methods affect the movement metrics that we collect and analyse, and on which we base our understanding of the behaviour in a population. Although, the effects of devices on animals are difficult to measure, we need to increase our efforts towards quantifying and understanding them. By underestimating the effects of devices we risk to invalidate the efforts of our research community in studying and preserving animal behaviour, especially in the case of threatened species which are often subjects of movement ecology studies.

The weight of a device, relative to an animal’s body weight (a currently accepted upper limit of to date 3%), is in general the only aspect currently considered in movement ecology studies involving tracking technology or marking methods [11]. Yet, also other aspects, such as device attachment (where and how the device is attached to the animal’s body) and device-induced drag, have been shown to affect flight behaviour [12]. In order to avoid these negative effects, researchers recommended to minimise the weight and flatten the shape of the devices to reduce drag [5, 9], to use flexible harnesses to reduce injuries for the birds [4, 13], and to position the devices at the centre of gravity to avoid destabilisation [1, 10, 14].

Patagial tags have been extensively used as markers in the last decades and have played an important role in behavioural studies. However, several studies confirmed adverse effects of patagial tags related to behaviour, nesting success and survival [15–17]. Studies on vultures, suggested that patagial tags might affect the aerodynamics of flight by impacting lift [18], which could negatively affect the energetic cost of flight. The severity of these effects also depends on tag material, as plastic tags might lift off the surface of the wing during flight and have a stronger impact compared to softer vinyl tags, which lie flat on the wing [19].

If the weight and drag of a tag on the wing were problematic *per se*, the way in which anatomic structures of the wing interact during flight makes the exact placement of such a tag even more so. A recent review on proper patagial tag placement highlighted how an incorrect placement of a tag on the patagium can cause injuries and result in the grounding of vultures [20]. Most of the vulture populations currently studied are endangered and in dire need of informed conservation action; hence studying their movements while finding the least invasive tagging technique to study their behaviour has become a priority. Researchers recently introduced leg bands as an alternative to patagial tags for individual identification [20]. In addition to marking methods, researchers use GPS tracking devices to follow individual trajectories and asses habitat use, resource selection and flight performance.

In this study, we compare the impact of patagial tags *vs* leg bands on the flight performance of an endangered soaring bird species, the Cape Vulture (*Gyps coprotheres*). Specifically, we investigate the effects of tag attachment in wild and captive bred individuals on flight probability, proportion of time spent flying, cumulative distance travelled and flight speed.

## Results

Between 2015 and 2019 we used GPS devices to track 27 vultures (14 wild and 13 captive bred) marked them with either patagial tags (15 birds) or leg bands (13 birds). The trajectories of the 27 individuals covered an area between −17° and −32° latitude. Individuals equipped with different tag attachments showed clear differences in the area they covered during the tracking period (figure 1). The final data set consisted of 5822 tracking days between April 2015 and November 2019, of which 2292 corresponded to birds equipped with leg bands (1079 of wild and 1213 of captive individuals), and 3530 to birds wearing patagial tags (1870 of wild and 1660 of captive individuals). The average tracking time for the individuals in the study was 2380.1 ± 302.7 h [mean ± SE; min = 172.5 h, max = 7027.6 h]. The number of segments per day considered in the analysis ranged from 1 to 82 and among all individuals, we recorded 2974 days (51%) in which no flight segment was recorded.

**Figure 1:**
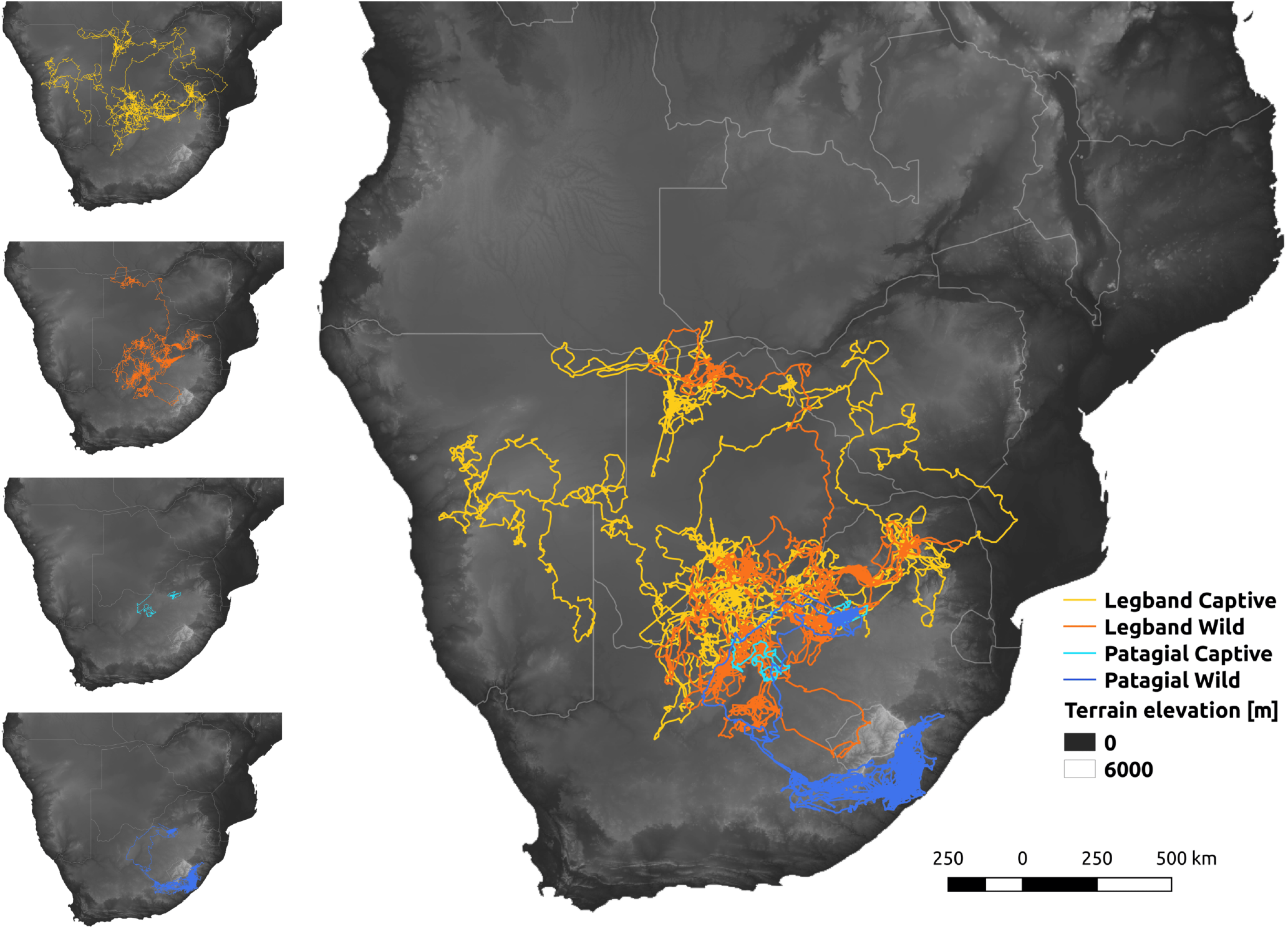
GPS trajectories of 27 African Cape Vultures, wild and captive, equipped with either leg bands or patagial tags. Colours show the four combinations of these categories.

In days when at least one flight segment was recorded, the birds flew only 11.3 ± 0.002 % [mean ± SE] of their daily tracking time, and covered a daily distance of 70.8 ± 1.2 km flying on average with a daily median ground speed of 6.9 ± 0.05 ms^−1^. A comparison of the raw data between attachment types and groups showed that the proportion of days with flight and the daily values of proportion of time spent flying, cumulative distance travelled and median flight speed were the lowest in captive birds equipped with patagial tags and the highest in birds wearing leg bands (similarly so between captive and wild; figure 2).

**Figure 2:**
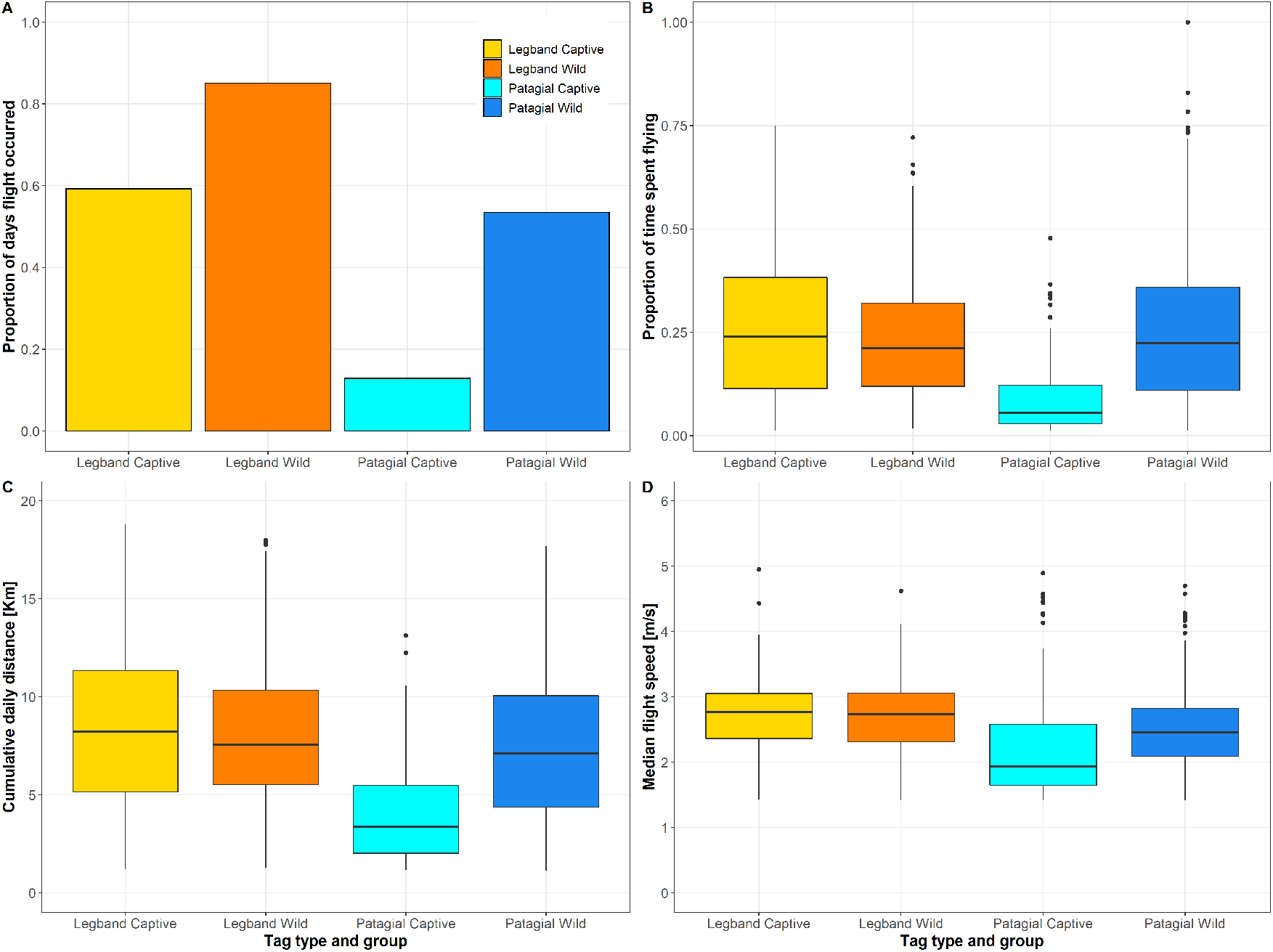
(A) Proportion of tracking days in which flight occurred, (B) daily proportion of time spent flying, (C) daily cumulative distance travelled, and (D) daily median flight speed, across groups. Different colours differentiate the four groups: captive bred individuals equipped with leg bands, wild individuals equipped with leg bands, captive individuals equipped with patagial tags and wild individuals equipped with patagial tags.

During the study, one captive vulture originally released with a patagial tag was found grounded, whereupon its patagial tag was replaced with a leg band (see Methods). After the tag replacement, we detected an increase in the flight parameters (figure 3) for this individual.

**Figure 3:**
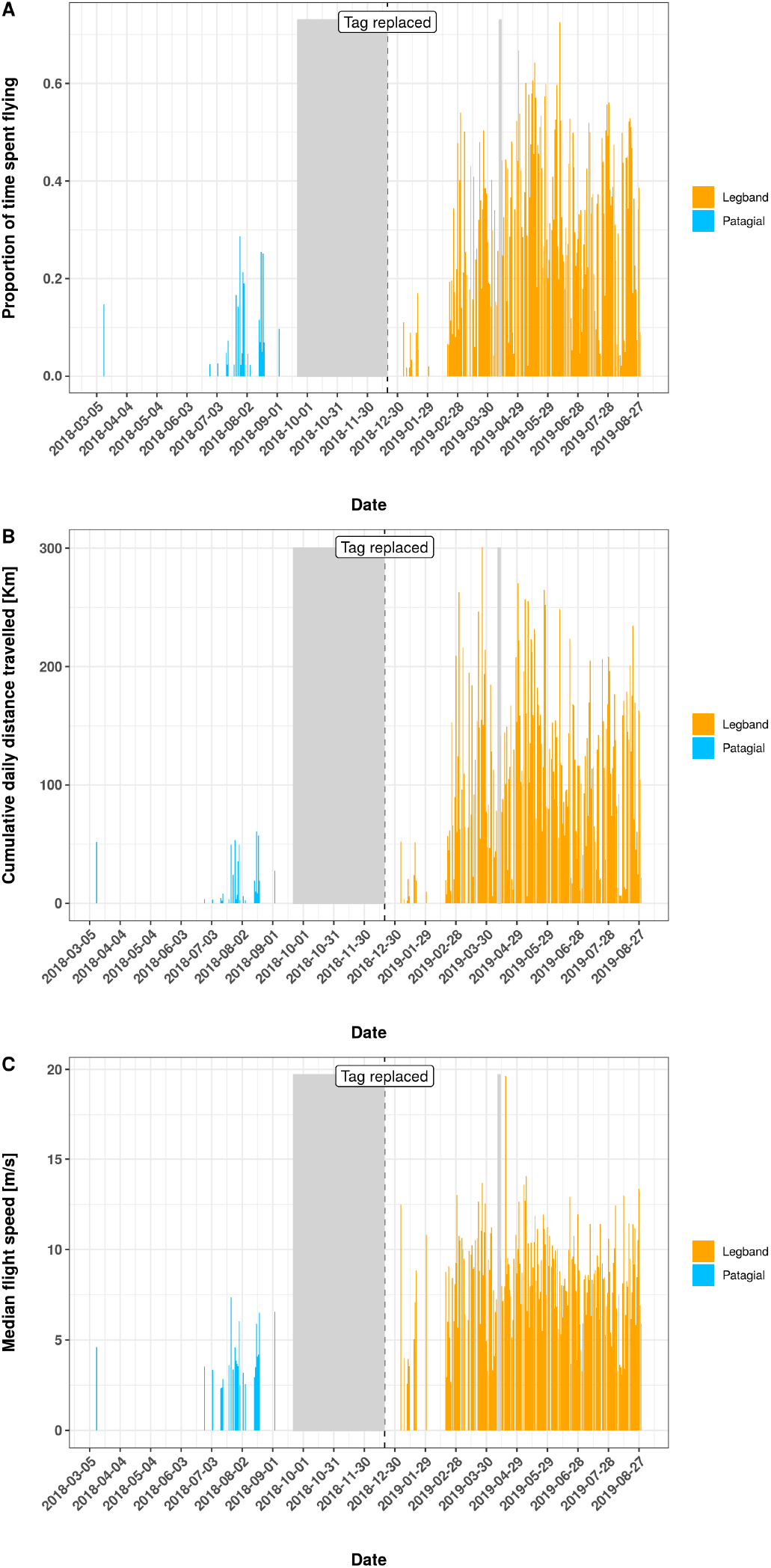
Increase in (A) daily proportion of time spent flying, (B) daily cumulative distance travelled, and (C) daily median flight speed for one individual vulture after replacing the patagial tag with a leg band. Days with variable values = 0 correspond to days in which no flight occurred. Gray polygons highlight days for which no tracking data were available.

Inferences about the effect of tag attachment and group on the flight performance of the birds were drawn based on the results of four generalized additive mixed models (GAMMs). Flight probability, proportion of time spent flying, daily cumulative distance travelled and flight speed were included as response variables. In all four models, attachment type and group (captive or wild) and their interaction term were included as predictors. In order to account for the uneven tracking effort we additionally included the number of daily locations and number of days since release.

### Occurrence of flight (flight probability)

Occurrence of flight (0 – absence, 1 – presence) was included as a response variable in the first GAMM. This model contained 5822 observations (days) of which 2292 belonged to the leg band group (0 = 656, 1 = 1636) and 3530 to the patagial group (0 = 2318, 1 = 1212), for a total of 27 individuals. The proportion of days in which no flight was recorded was higher for individuals wearing patagial tags (65.6 %) compared to those wearing leg bands (28.6 %; figure 2). Our model predicted flight probability to be lower (albeit not significantly) for wild individuals equipped with patagial tags compared to wild individuals equipped with leg bands (patagial = −1.47 ± 0.79 [estimate ± SE]). The effect of patagial tags on flight probability was the strongest and lower than the significance threshold (p<0.05) for captive individuals equipped with patagial tags (patagial:captive = −2.20 ± 0.83). We detected no significant difference between captive and wild individuals equipped with leg bands. Indeed, our model predicted the highest probability to fly (84 %) for wild individuals with leg bands and the lowest (6.47 %) for captive individuals with patagial tags. Among-individual variability was quite high, with 95 % of the individuals having a flight probability between 24.5 % and 98.8 %. However, the variability attributable to individual variation (intercept SD = 1.42) was lower than the effect associated with the patagial attachment both on wild and captive individuals (table 1).

**Table 1:**
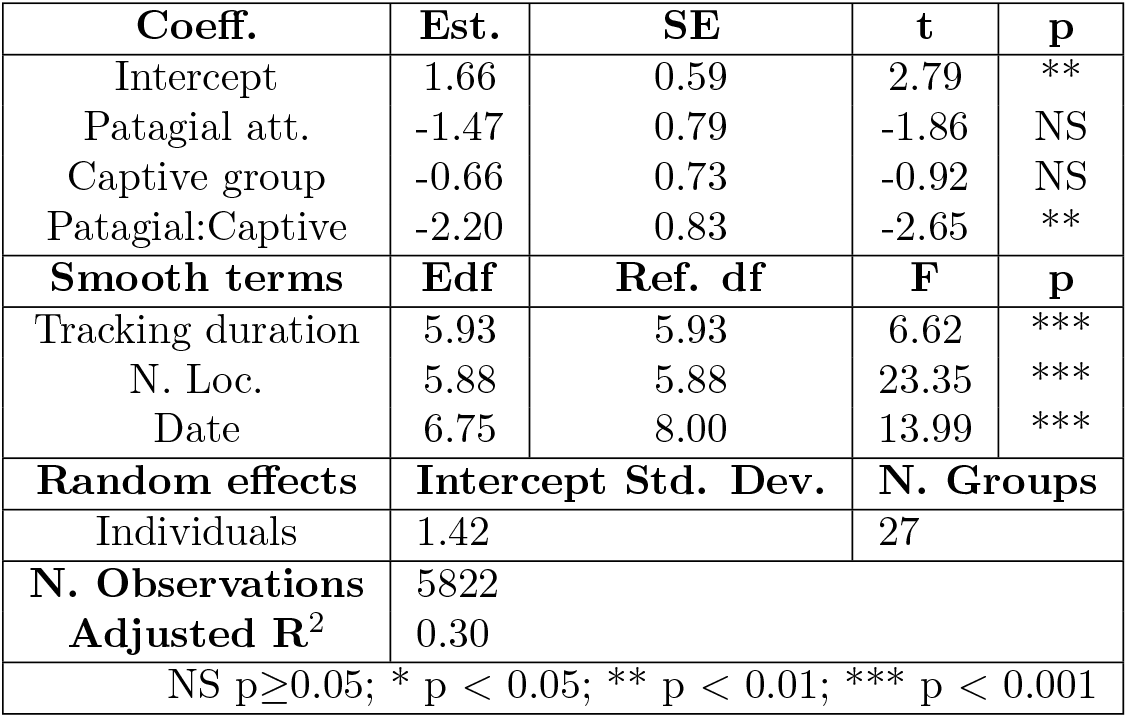
GAMM with occurrence of flight included as dependent variable, interaction between tag attachment and group as fixed term and individual identity as random intercept. Number of days since deployment, number of locations and day of the year were included as smooth terms. The model was fitted with the restricted maximum likelihood using the binomial family and a logit link function.

| Coeff. | Est. | SE | t | p |
| --- | --- | --- | --- | --- |
| Intercept | 1.66 | 0.59 | 2.79 | ** |
| Patagial att. | -1.47 | 0.79 | -1.86 | NS |
| Captive group | -0.66 | 0.73 | -0.92 | NS |
| Patagial:Captive | -2.20 | 0.83 | -2.65 | ** |
| Smooth terms | Edf | Ref. df | F | p |
| Tracking duration | 5.93 | 5.93 | 6.62 | *** |
| N. Loc. | 5.88 | 5.88 | 23.35 | *** |
| Date | 6.75 | 8.00 | 13.99 | *** |
| Random effects | Intercept Std. Dev. |  | N. Groups |  |
| Individuals | 1.42 |  | 27 |  |
| N. Observations | 5822 |  |  |  |
| Adjusted R<sup>2</sup> | 0.30 |  |  |  |
| NS p≥0.05; * p < 0.05; ** p < 0.01; *** p < 0.001 |  |  |  |  |

All smooth terms included in the model – the duration, i.e. the number of days since deployment, number of locations and the Julian date – had a significant effect on the occurrence of flight. Flight probability increased with the number of locations until around 50 locations, indicating that using a number of daily segments < 50 could result in a lower probability of detecting flight. Flight probability was highest (> 50%) between the end of May and the beginning of September (days 150 to 250). The tracking duration (number of days since deployment) positively influenced flight probability after 500 days (about 1.4 years) from deployment (see figure 1 in Additional file 1).

The following three models were all run only on days in which at least one flight segment was recorded (n = 2848 days, 1636 corresponding to leg band and 1212 to the patagial group for 27 individuals.

### Daily proportion of time spent flying

Daily proportion of time spent flying did not differ significantly for individuals wearing patagial tags, compared to leg bands, nor did it differ between captive and wild individuals. The difference between the groups spending the highest and the lowest proportion of their time flying was rather small: 0.20 for wild individuals wearing patagial tags (patagial = 0.11 ± 0.14) and 0.16 for captive individuals with patagial tags (patagial:captive = −0.17 ± 0.14). 95 % of the individuals showed a proportion ranging from 0.11 to 0.27, making among-individuals variability high (intercept SD = 0.26) compared to the effect of tag attachment and group (table 2).

**Table 2:** GAMM with daily proportion of time spent flying included as dependent variable, interaction between tag attachment and group as fixed term and individual identity as random intercept. Number of days since deployment, number of locations and day of the year were included as smooth terms. The model was fitted with the restricted maximum likelihood using the binomial family and a logit link function.

| Coeff. | Est. | SE | t | p |
| --- | --- | --- | --- | --- |
| Intercept | -1.51 | 0.11 | -14.23 | *** |
| Patagial att. | 0.11 | 0.14 | 0.77 | NS |
| Captive group | -0.09 | 0.13 | -0.69 | NS |
| Patagial:Captive | -0.17 | 0.14 | -1.20 | NS |
| Smooth terms | Edf | Ref. df | F | p |
| Tracking duration | 7.28 | 7.28 | 26.03 | *** |
| N. Loc. | 8.92 | 8.92 | 16584.51 | *** |
| Date | 7.68 | 8.00 | 210.37 | *** |
| Random effects | Intercept Std. Dev. |  | N. Groups |  |
| Individuals | 0.26 |  | 27 |  |
| N. Observations | 2848 |  |  |  |
| Adjusted R<sup>2</sup> | 0.93 |  |  |  |
| NS p≥0.05; * p < 0.05; ** p < 0.01; *** p < 0.001 |  |  |  |  |

Tracking duration, number of daily locations and Julian date, all included as smooth terms in the model, had a statistically significant effect on the proportion of time spent flying. Proportion of time spent flying increased with the number of daily locations and reached its maximum at around 30 locations, indicating that recording < 30 segments per day would decrease our chances to identify flight segments. The model showed a small seasonal effect on the proportion of time spent flying, with a small increase between the beginning of April and the end of October. The proportion of time spent flying did not to be influenced by the duration of the tracking period (see figure 2 in Additional file 1).

### Daily distance travelled

Wild individuals equipped with patagial tags travelled significantly shorter distances, of about 10 km less, compared to wild individuals equipped with leg bands (effect size patagial = −0.65 ± 0.22). In captive bred individuals, the difference between distance travelled when wearing leg bands and patagial tags was smaller than in wild individuals (about 7 km) and statistically non-significant. When comparing within the same tag attachment type, we did not find any significant difference between wild and captive bred individuals. The random structure of the model showed small among-individual variation, with 95 % of the individuals travelling between 50 and 72 km/day. The variability attributable to individual variation (intercept SD = 0.37) was lower relative to the effect associated to the patagial attachment on wild birds (table 3).

**Table 3:** GAMM with the square root of the daily cumulative distance included as dependent variable, interaction between tag attachment and group as fixed term and individual identity as random intercept. Number of days since deployment, number of locations and day of the year were included as smooth terms. The model was fitted with the restricted maximum likelihood using the gaussian family.

| Coeff. | Est. | SE | t | p |
| --- | --- | --- | --- | --- |
| Intercept | 7.79 | 0.16 | 49.42 | *** |
| Patagial att. | -0.65 | 0.22 | -2.97 | ** |
| Captive group | -0.09 | 0.21 | -0.44 | NS |
| Patagial:Captive | 0.19 | 0.27 | 0.70 | NS |
| Smooth terms | Edf | Ref. df | F | p |
| Tracking duration | 5.34 | 5.34 | 6.37 | *** |
| N. Loc. | 7.28 | 7.28 | 3568.63 | *** |
| Date | 5.45 | 8.00 | 3.76 | *** |
| Random effects | Intercept Std. Dev. |  | N. Groups |  |
| Individuals | 0.37 |  | 27 |  |
| N. Observations | 2848 |  |  |  |
| Adjusted R<sup>2</sup> | 0.92 |  |  |  |
| NS p≥0.05; * p < 0.05; ** p < 0.01; *** p < 0.001 |  |  |  |  |

The positive and significant effect of the number of daily locations suggests that collecting a higher number of segments per day would result in recording larger daily distances. Our model showed no seasonal effect on the distance travelled and no increase or decrease in daily movement with the number of days from release (see figure 3 in Additional file 1).

### Daily median flight speed

Wild individuals equipped with patagial tags reached significantly lower flight speeds (about 0.88 ms^−1^ less) compared to wild individuals wearing leg bands (effect size patagial = −0.17 ± 0.07). The model showed no significant differences between wild and captive bred individuals, however the effect sizes suggest that captive individuals wearing leg bands travelled at the fastest speeds, about 7.2 ms^−1^, while captive individuals with patagial tags at the lowest, about 5.9 ms^−1^ (patagial:captive = −0.09 ± 0.09). Among-individuals variability was relatively high (intercept SD = 0.10) if compared to the largest effect size due to the patagial attachment, and 95 % of the individuals showed a daily median speed between 6 and 8 ms^−1^ (table 4).

**Table 4:** GAMM with the square root of the daily median flight speed included as dependent variable, interaction between tag attachment and group as fixed term and individual identity as random intercept. Number of days since deployment, number of locations and day of the year were included as smooth terms. The model was fitted with the restricted maximum likelihood using the gaussian family.

| Coeff. | Est. | SE | t | p |
| --- | --- | --- | --- | --- |
| Intercept | 2.67 | 0.05 | 58.34 | *** |
| Patagial att. | -0.17 | 0.07 | -2.59 | ** |
| Captive group | 0.02 | 0.06 | 0.36 | NS |
| Patagial:Captive | -0.09 | 0.09 | -1.05 | NS |
| Smooth terms | Edf | Ref. df | F | p |
| Tracking duration | 5.60 | 5.60 | 7.20 | *** |
| N. Loc. | 6.30 | 6.30 | 85.68 | *** |
| Date | 4.50 | 8.00 | 3.06 | *** |
| Random effects | Intercept Std. Dev. |  | N. Groups |  |
| Individuals | 0.10 |  | 27 |  |
| N. Observations | 2848 |  |  |  |
| Adjusted R<sup>2</sup> | 0.26 |  |  |  |
| NS p≥0.05; * p < 0.05; ** p < 0.01; *** p < 0.001 |  |  |  |  |

The number of daily locations, duration of tracking and Julian date significantly influenced median flight speed. Daily median flight speed significantly increased with the number of daily locations and reached an asymptote at around 25 locations. Flight speed also increased until around 200 days (about seven months) after deployment but this increase was not consistent over time. The model also showed a seasonal effect on the daily flight speed, with a slight decrease in speed between the beginning of April and the end of October (see figure 4 in Additional file 1).

## Discussion

We have demonstrated that tag attachment has a severe impact on the flight performance of Cape Vultures. Our results showed that both wild and captive individuals equipped with patagial tags were significantly less likely to take flight (lower proportion of days in which flight occurred) and when doing so flew at significnatly lower ground speed compared to individuals wearing leg bands. Although statistically not significant, we believe it to be relevant that among all groups, captive individuals wearing patagial tags also spent the least proportion of time flying. Finally and as a consequence of the above, our models also showed that wild birds equipped with patagial tags travelled significantly shorter distances compared to those wearing leg bands.

Age and experience are known to affect ranging behaviour and distance travelled in vultures, with younger individuals moving more than older ones, who already established their breeding territories [21, 22]. In this study, due to our small sample size, it was not possible to account for age in the models. Our dataset included 21 individuals among which were fledglings, juveniles and subadults and only six adults. However, the adult individuals were equally distributed between the two attachment types, thus we are confident that omitting age from the models did not bias our findings.

Shorter daily distance travelled means smaller area covered every day by the birds to forage. Vultures are scavengers and as such cover large areas to feed on ephemeral food resources [23]. At individual and population level, a restricted flight potential and therefore a restricted area available to forage might not be a problem if food resources are abundant, but they surely reduce an individual’s ability to react to changes in food availability and environmental conditions. A restricted area covered daily by these birds can also lead to ecosystem-level effects. Scavengers play an important role in the ecosystem thanks to the services they provide, such as preventing the spread of infectious diseases, recycling organic material into nutrients and stabilising food webs [24]. Therefore a restricted flight potential and reduction in the area covered by these birds caused by improper tag attachment can have far-reaching consequences at the ecosystem level.

Our results are consistent with previous studies, showing that tag attachment influences foraging trip duration. In some studies, birds carrying extra weight had shorter foraging trips [10, 14]. Other studies investigating the effect of device weight however showed that birds carrying devices exhibited longer foraging trips [6, 25, 26], suggesting that individuals wearing extra weight might have to compensate for their increased greater energy demands by travelling more and farther in search for food [12, 14, 27, 28]. This might raise questions about whether the decrease in the four flight parameters included in our study should be considered detrimental for the birds.

Unfortunately, we did not have access to information about body condition, fitness or survival rate of the individuals after they were tagged. However, birds equipped with patagial tags were more often found grounded, injured or with incorrect tag placement compared to birds wearing leg bands ([20], K. Wolter pers. obs.). In addition, birds in our study were equipped with the same type of tracking device and therefore experienced the same additional weight, the only difference was the marking method. Previous studies suggested that in vultures, a device attached on the wing can influence the aerodynamics of flight [18, 19], while to our knowledge no negative effects on flight performance have been reported for devices attached to the leg. Our suggestion of an adverse effect of patagial tags is further supported by the anecdotal evidence of one individual vulture sequentially equipped with both types of attachment. This captive vulture, originally equipped with a patagial tag and released, was found grounded about 6 months later and after rehabilitation its tag was replaced with a leg band. The data showed that after the tag replacement, all flight parameters measured for this individual increased to match the movement parameters of the other leg banded individuals, suggesting that this individual was rather hampered previously in its movement.

More research is needed to fully understand the consequences of different tagging methods on vultures, including their direct effects on body condition, reproduction and survival. Nonetheless, we urgently recommend using leg bands instead of patagial tags. Should the use of patagial tags be considered unavoidable, we urge researchers to consult experts about the exact positioning of the tag on the patagium (see the manual in [20]). Some researchers in South Africa already started replacing patagial tagging with leg banding. Fast implementation of such changes, in response to up cutting-edge research on the topic, are especially important for endangered and critically endangered species, such as the Cape Vulture and other African vulture species which are declining in numbers dramatically [29]. In such species, the use of the least invasive tagging method can have a significant impact on their population stability and extinction risk.

Tracking devices and tagging methods are under constant development and the body of literature assessing their impact is inevitably not comprehensive. The challenging task for the movement ecology research community is to keep up with the methodological developments by studying how different devices and attachments affect animal behaviour in order to improve current methods to minimise their impact [13]. Equipping birds with devices generally affects many aspects of their behaviour and ecology [4]. In several cases, however, studies reported contradicting results of the effects of different devices on birds [4, 30]. For example, some studies showed adverse effects of patagial tags while others reported no effects [30]. The reason for that could be that different species respond differently to the attachment of devices because of their ecological differences [6]. When assessing which device to use, it is therefore necessary to understand its impacts on the specific study species or on phylogenetically related or functionally similar species. Discrepancies between studies may also be generated by differences in the temporal and special scales at which data were collected, or by differences in the environmental conditions the animals experienced [6]. Therefore, to gain a better understanding of the impact of tagging methods, it is important to take into account possible differences due to methodology (e.g. tag type, sampling schedule), environmental context (e.g. seasonality), and different aspects of the species’ biology (i.e. flight, body condition, reproduction) [4, 6].

## Conclusions

Researchers need to balance the benefits of data acquired by tracking animals with the adverse consequences on the animals’ health and the potential biases on the scientific results. These biases invalidate the comparability of information across individuals and populations tagged with different methods, and defeat the purpose of generalising our behavioural conclusions beyond the single device and attachment. As movement ecologists we aim to study the movement behaviour of animals in different conditions to predict how different species or population would react to changes in their environment. Comparisons at different spatial and temporal scales are therefore an important part of our field of research, but these comparisons risk to become meaningless if we choose to ignore the biases due to how we measure the phenomenon we observe.

## Methods

### Data collection

Between 2015 and 2019, we tracked 27 Cape Vultures in South Africa (14 males, 13 females of which six were adults, four fledglings, 16 juveniles and one was a subadult). Thirteen vultures were captive bred and 14 wild. We equipped individuals with backpack mounted GPS devices and marked them with either a leg band (13 birds) or patagial tags (15 birds). Patagial tags were placed on both wings, while leg band was fitted only on one leg. Age classes by tag type and group are presented in Additional file 1, table 1. One individual was released twice, wherefore there are 28 tracking events of 27 individuals included in the study. This captive bred bird was first released on March 5, 2018 and equipped with GPS and patagial tag. On September 21, the same bird was found grounded and showed no interest in flying. Its patagial tag was removed and the bird was rehabilitated. On December 27, 2018 the bird was equipped with GPS and leg band and released back into the wild. This allowed us to evaluate the effect of the tag replacement on the bird’s flight performance.

For tracking, we used second and third generation GPS-GSM Wildlife Telemetry System from Cellular Tracking Technologies (1000-BT3 and 1000a series). The devices sampled positions 24 h/day at the median of either 1, 5 or 15 min. Sampling frequency for some individuals was reduced at night, with 90 % of locations across all individuals being collected between 4:00 and 16:00. The accuracy of the GPS devices was given at 2.5 m (50% CEP statistical test) or 5 m (95 % 2dRMS statistical test).

Captive birds were born and raised in captivity at VulPro’s rehabilitation centre and released before the age of one. All wild birds were found grounded due to minor injuries (e.g. heat stroke, starvation, weather) and admitted to the rehabilitation centre at VulPro for a couple of months before being released back into the wild. Before 2018, birds were released from the VulPro rehabilitation centre (captive-bred = 2, wild = 6) whereas after 2018, from the release site located at Nooitgedacht (captive-bred = 11, wild = 8). Both captive and wild birds were equipped upon release with GPS devices and tagged with either legband or patagial tags. Capturing, handling, fitting tracking devices and patagial tags on vultures was done following protocols described in [31].

### Data processing

The initial data set included a total of 625,861 GPS locations from 27 individuals and 28 tagging events. We manually removed GPS locations falling outside the African continent. We then removed additional outliers based on speed between consecutive locations using the R package ctmm to account for location error [32]: we set a conservative GPS error (user equivalent range error) of 10 m, used the functions available in the ctmm package to compute speed and variance in speed at each location, and repeatedly removed locations with speed > 25 ms^−1^ [33]. This procedure was only used to detect and remove outliers but not for statistical purposes. The map in figure 1 shows the distribution of the clean trajectories.

After removing outliers we thinned the trajectories, in order to minimise potential biases in speed and distance calculations due to variable sampling frequency. Thinning was done by selecting from the original track only segments with 15±5 min time lag. We then calculated time lag, horizontal displacement and speed between locations of the selected segments. Both thinning and variable calculations were performed using the R package move [34].

For the next part of the analysis, we excluded segments with variance in speed > 0.5 ms^−1^ and time lag > 1 hour because we considered them less reliable in the speed calculation. We classified the remaining segments into flying and not-flying based on a speed threshold of 2 ms^−1^ which we used to calculate metrics of flight performance to evaluate the effects of tag attachment. Specifically, for each individual and day of tracking we calculated: (i) Occurrence of flight (1 if at least one flight segment was recorded in the day, 0 otherwise); (ii) Proportion of time spent flying in a day(sum of flight time divided by tracking time for each day); (iii) Cumulative distance travelled (sum of horizontal displacements during flight segments, in km); and (iv) Median flight speed (median of the speeds achieved during flight segments, in ms^−1^).

### Data analysis

We used four GAMMs (R package mgcv [35] to test the effect of tag attachment on each of the above flight parameters. In all four models, attachment type and group (captive or wild) and their interaction term were included as fixed terms. The number of locations and number of days since release (deployment date) were also included in the models to account for the uneven tracking effort (number of locations and tracking duration) between days and individuals. These two parameters were included in the model as smooth terms (thin plate regression splines). Given the year-round tracking, we additionally included in the model Julian date (day of the year in the range 1-366) as cyclic cubic regression spline, to account for non-linear effects of seasonality. Finally, individual identity was included as a random effect in all models. Models including the proportion of time spent flying and occurrence of flight as response variables were fitted as logistic regressions with a “logit” link function. The variables cumulative distance travelled and median flight speed were square-root transformed and the corresponding models fitted using a Gaussian error distribution. Data processing and analysis were performed in R [36]. The complete R scripts to reproduce the analysis can be found in Additional file 2.

### Ethics approval and consent to participate

Work with Cape Vultures in this study was performed in accordance with ethical legislation (permits in Gauteng: 0-002619, North West Province: 10846 and 5723, and Limpopo: 06758 and 13677).

## Acknowledgements

We would like to thank Alexandra Howard for her contribution in data preparation. We would also like to thank the landowners, farmers and volunteers who have assisted us with vulture resightings, vulture collections and rehabilitation needs.

## Additional Files

- Additional file 1 — “AdditionalFile1 _suppl.pdf”: File containing supplementary table and figures
- Additional file 2 — “AdditionalFile2_Rcode.pdf”: File containing the complete R code to reproduce the data processing and analysis.

## Table abbreviations

Coeff.: coefficient
Est.: estimate
Edf: estimated degrees of freedom
Ref.df: reference degrees of freedom
Std. Dev: standard deviation
N.: number
NS: not significant

