## Additional file 1 - Supplementary material for "Wing tags severely impair movement in African Cape Vultures"

Table 1: Age classes by tag type and group.

| Tag type | Group | Age | N of ind |
|---|---|---|---|
| Legband | Captive | Adult | 0 |
| Patagial | Captive | Adult | 0 |
| Legband | Wild | Adult | 2 |
| Patagial | Wild | Adult | 4 |
| Legband | Captive | Fledgling | 0 |
| Patagial | Captive | Fledgling | 0 |
| Legband | Wild | Fledgling | 3 |
| Patagial | Wild | Fledgling | 1 |
| Legband | Captive | Juvenile | 7 |
| Patagial | Captive | Juvenile | 7 |
| Legband | Wild | Juvenile | 1 |
| Patagial | Wild | Juvenile | 2 |
| Legband | Captive | Subadult | 0 |
| Patagial | Captive | Subadult | 0 |
| Legband | Wild | Subadult | 0 |
| Patagial | Wild | Subadult | 1 |

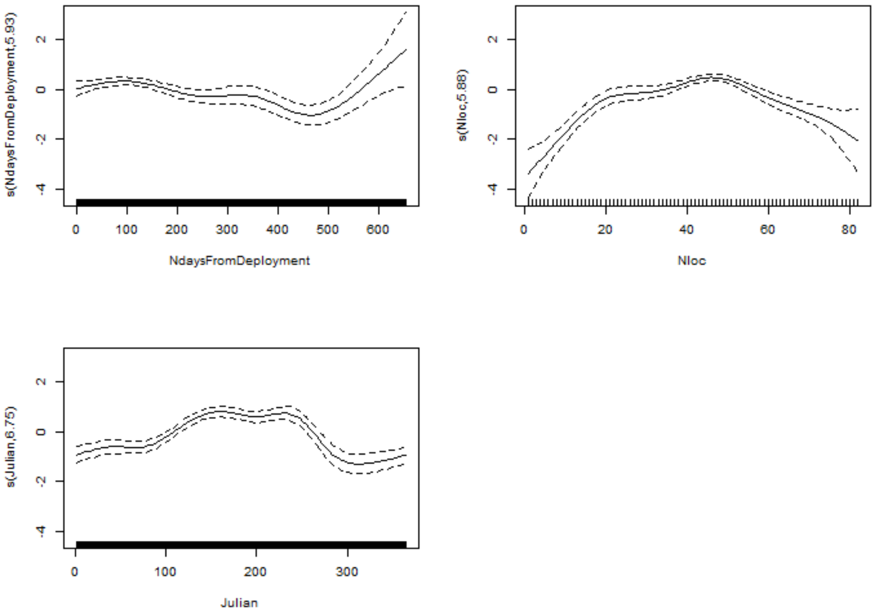


Figure 1: The occurrence of flight in relation to the three smooth terms included in the model: (A) number of days since deployment, (B) number of locations, and (C) Julian date.

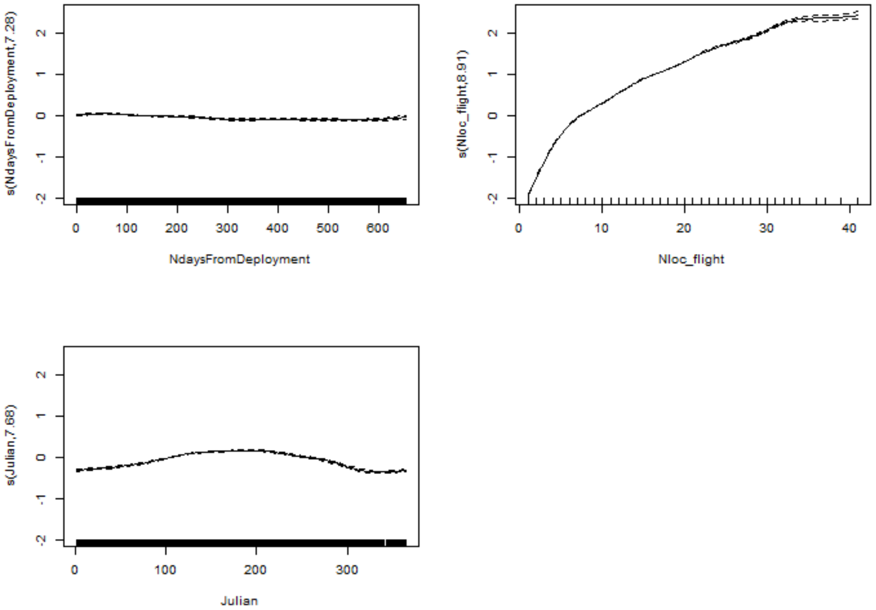


Figure 2: The proportion of time spend flying in relation to the three smooth terms included in the model: (A) number of days since deployment, (B) number of locations, and (C) Julian date.

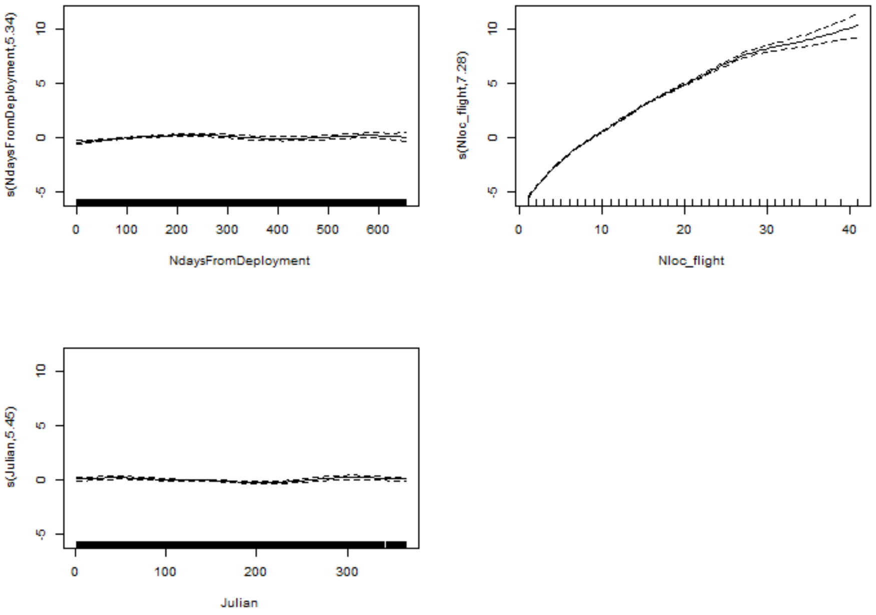


Figure 3: Cumulative daily distance travelled in relation to the three smooth terms included in the model: (A) number of days since deployment, (B) number of locations, and (C) Julian date.

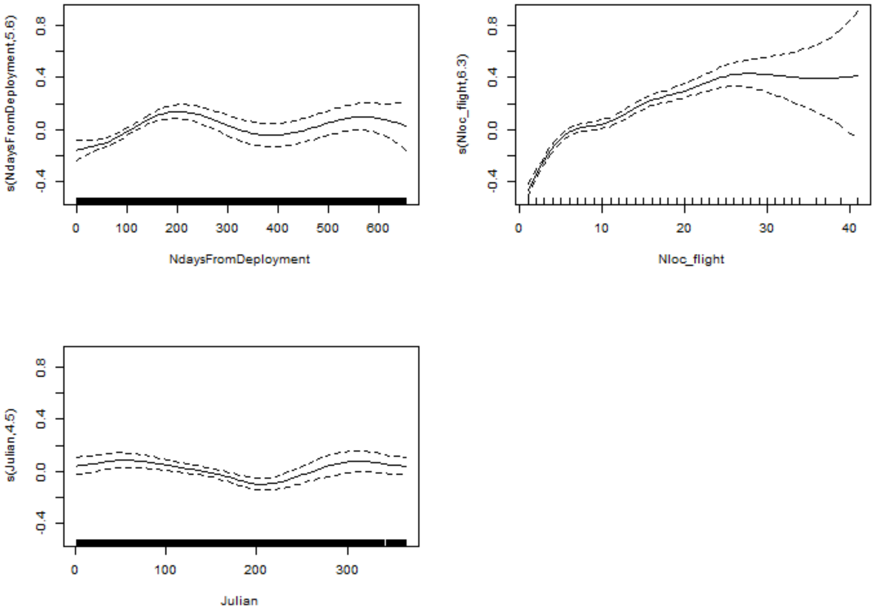


Figure 4: Median flight speed in relation to the three smooth terms included in the model: (A) number of days since deployment, (B) number of locations, and (C) Julian date.
