## Additional file 2 - R code for "Wing tags severely impair movement in African Cape Vultures"

Data processing and analysis were performed in R. We divided the procedure in different steps. An output file is produced at the end of each step.

### Step 1 - Combining individual datasets

```
path <- "/home/mscacco/"

setwd(paste0(path,"ownCloud/VulturesMPI-VulProKerry"))

# Data import, match columns, add ID ####
#_________________________________________
# Import data and fill in empty ID data (if ID column already exist, in folder "allBirds")
##
list1 <- list.files("allBirds", pattern="csv", full.names=T)
list_files1 <- lapply(list1, read.csv, na.strings = c("NA",""), as.is=T)
# Fill in empty ID rows
list_files1 <- lapply(list_files1, function(x){
  x$ID <- unique(x$ID)[!is.na(unique(x$ID))]
  return(x)})
# rbind all data from the first batch of individuals
df1 <- do.call(rbind,list_files1)
head(df1)
tail(df1)
length(unique(df1$ID))

# Import data and add column ID if not existing (folder "allBirds/Additional_IDs")
##
list2 <- list.files("allBirds/Additional_IDs", pattern="csv", full.names=T)
list_files2 <- lapply(list2, read.csv, na.strings = c("NA",""), as.is=T)
# Element 8 has different columns
lapply(list_files2, head)
# Try to make dataset 8 match the column names of the others
list_files2_8 <- as.data.frame(matrix(NA, nrow=nrow(list_files2[[8]]),
                                      ncol=ncol(list_files2[[1]])))
names(list_files2_8) <- names((list_files2[[1]]))
list_files2_8[,c("GPS_date_YYYY.MM.DD","GPS_utc_HH.MM.SS","GPS_YYYY.MM.DD_HH.MM.SS")] <-
  cbind(substr(list_files2[[8]]$timestamp,1,10),
        substr(list_files2[[8]]$timestamp,12,19),
        substr(list_files2[[8]]$timestamp,1,19))
list_files2_8[,c("lon","lat","hdop","speed","alt","data_voltage")] <-
  list_files2[[8]][,c("location.long","location.lat","gps.hdop","ground.speed",
                    "height.above.msl","tag.voltage")]
head(list_files2_8)
# And element 8 in the list with the new one
list_files2[[8]] <- list_files2_8
```

```
# Extract IDs from filename
IDs <- sapply(strsplit(list2, " "), "[", 2)
# Now add the column ID to each dataframe and re-order the columns as in df1
list_files2_id <- mapply(x=list_files2, y=IDs, function(x,y){
  x$ID <- y
  x <- x[,names(df1)] #order columns as in df1
  return(x)
}, SIMPLIFY = F)
# Now that they match rbind all data from the additional individuals
df2 <- do.call(rbind,list_files2_id)

# Now that they contain the same columsn we can rbind df1 and df2
##
table(names(df1)%in%names(df2))
length(unique(df1$ID)) + length(unique(df2$ID))

df <- rbind(df1, df2)
length(unique(df$ID)) # we have 39 unique tracking periods
anyNA(cbind(df$lon,df$lat))
nrow(df[which(is.na(df$lat)|is.na(df$lon)),])
df <- df[which(!is.na(df$lat)|!is.na(df$lon)),]

# Merge extra infos, correct timestamps ####
#___________________________________________

# Add extra infos (additional variables) into the dataset
extra_info <- read.csv("extra_info_SexAge.csv", row.names=NULL,
                       na.strings = c("NA",""), as.is=T)
table(extra_info$Tracker.ID)
#individual G34918 had patagial (wing) for some time and then replaced with legband
table(extra_info$Animal.ID)
length(unique(extra_info$Tracker.ID)) #38 unique tracking periods
length(unique(extra_info$Animal.ID)) #on 37 individuals - 1 individual (G34918) tagged twice
head(df)
head(extra_info)

summary(unique(df$ID) %in% unique(extra_info$Tracker.ID))

# Merge both dataset by ID
df_compl <- merge(df, extra_info, by.x="ID", by.y="Tracker.ID", all.x=T)

nrow(df_compl) == nrow(df)
length(unique(df_compl$ID))
length(unique(df_compl$Animal.ID))
anyNA(df_compl$Animal.ID)

#Change timestamp format
library(lubridate)
df_compl$dates <- parse_date_time(df_compl$GPS_date_YYYY.MM.DD,
                                  orders = c('ymd', 'dmy', 'mdy')) # Uniform date formats
df_compl$timestamp <- as.POSIXct(paste0(as.character(df_compl$dates)," ",
                                        df_compl$GPS_utc_HH.MM.SS),
                                 format="%Y-%m-%d %H:%M:%S", tz="UTC") #convert into posixct
```

```
anyNA(df_compl$timestamp) #check that works, no NAs in timestamps
anyNA(df_compl$dates)

# Remove datetime columns with errors
df_compl <- df_compl[, !names(df_compl)%in% c("GPS_date_YYYY.MM.DD", "GPS_utc_HH.MM.SS",
                                              "GPS_YYYY.MM.DD_HH.MM.SS", "CSV")]

# Save correct dataset
write.csv(df_compl,file="out1_vulture_dataset_allIndividuals.csv",row.names = FALSE)
save(df_compl, file="out1_vulture_dataset_allIndividuals.rdata")
```

### Step 2 - Data cleaning

```
path <- "/home/mscacco/"
#path <- "C:/Users/Martina.ska/Documents/Martina/"
setwd(paste0(path,"ownCloud/VulturesMPI-VulProKerry"))

# Load the vulture dataset
load("out1_vulture_dataset_allIndividuals.rdata") #object df_compl

# Remove duplicated timestamps ####
#__________________________________

# Order by ID and timestamps
df_compl <- df_compl[order(df_compl$Animal.ID, df_compl$ID, df_compl$timestamp),]

# Check that there is no NAs in timestamps, coordinates and IDs
anyNA(df_compl$timestamp)
anyNA(df_compl$lon)
anyNA(df_compl$lat)
anyNA(df_compl$Animal.ID)
anyNA(df_compl$ID)

# plot(df_compl$lon, df_compl$lat, pch=19)

# Check for duplicated timestamps, no duplicates!!!
dup <- getDuplicatedTimestamps(x=as.factor(df_compl$ID),
                               timestamps=df_compl$timestamp)
length(dup)

# Transform in movestack object ####
#___________________________________

library(move)

ms <- move(x=df_compl$lon,
           y=df_compl$lat,
           time=df_compl$timestamp,
           data=df_compl,
           proj=CRS("+proj=longlat +datum=WGS84"),
           animal=df_compl$ID) # Use column ID to identify separate tracking period
```

```
# Check tracking duration/hours of tracking per day ####
#______________________________________________________
library(lubridate)

hist(hour(df_compl$timestamp))
quantile(hour(df_compl$timestamp), seq(0,1,0.01))
mean(hour(df_compl$timestamp));sd(hour(df_compl$timestamp))
lapply(split(ms), function(m) range(timestamps(m)))
lapply(split(ms), function(m) table(hour(timestamps(m))))


# Remove outliers based on Longitude ####
#_______________________________________

# Plot on the map and see broad outliers
library(maptools)
data("wrld_simpl")
plot(crop(wrld_simpl, extent(ms)))
lines(ms)
# draw broad extent by hand and use it to crop the clear outliers
#maxExt <- drawExtent()
#save(maxExt, file="cropExtent.RData")
load("cropExtent.RData")
ms_long <- crop(ms, maxExt)

# Result
sp::plot(crop(wrld_simpl, extent(ms_long)))
lines(ms_long)

# Remove speed outliers using ctmm ####
#_____________________________________

library(parallel) #for detectCores()
library(doMC) #for parallel=T
library(plyr) #for llply
library(lubridate)
library(ctmm)
# vignette("error")
# vignette("variogram")

if(dir.exists("plotOutliers_ctmm")==F){dir.create("plotOutliers_ctmm")}

#registerDoMC(detectCores()-1)
ms_clean <- moveStack(llply(1:n.indiv(ms_long), .fun=function(i){
  print(paste0("Working on animal ",i," of ",n.indiv(ms_long)))
  m <- ms_long[[i]]
  id <- paste(i,m@idData$Species,m@idData$Tag.Type,m@idData$Animal.ID,m@idData$ID, sep="_")
  ## Convert into telemetry object
  t <- as.telemetry(m)
  ## Assign a conservative error value of 10 m (company said 2.5-5 m)
  uere(t) <- 10
  out <- outlie(t, plot=F)

  ## Iteratively remove outliers with speed > 25 m/s
```

```
  while(any(out$speed > 25)==T){
    t <- t[which(out$speed <= 25),]
    out <- outlie(t, plot=F) # plotting off during the loop
  }
  ## Associate columns with speed and variance from outlie to telemetry object
  colnames(out)[1:2] <- c("outlieSpeed","outlieDistance")
  tdata <- cbind(t, out)
  ## Associate the telemetry information with the idData information and create a data table
  tdata <- cbind(tdata, m@idData)
  ## Transform the t object in move and associate data from the datatable
  #to restore all the information from the original move object
  mv <- move(x=t$longitude, y=t$latitude,
             time= as.POSIXct(t$t,origin="1970-01-01",tz=t@info$timezone),
             proj= CRS("+proj=longlat +ellps=WGS84"),
             animal=tdata$ID,
             data=tdata)
  return(mv)
}, .parallel=F, .progress = "."), forceTz="UTC")

## save the move stack object cleaned from the outliers
save(ms_clean, file="out2_ctmm_vulture_movestack_noOutliers.rdata")


## Thin data so that all have a comparable timelag
#___________________________________________________

head(ms_clean)
allTimelags <- do.call(c,lapply(split(ms_clean), function(m)as.numeric(timeLag(m, units="mins"))))
summary(allTimelags)
hist(allTimelags, breaks="FD", xlim=c(0,60)) #most common is around 15 minutes
sort(table(allTimelags), decreasing = T) # most fixes around 1, 5 and 15 mins
sort(table(round(allTimelags)), decreasing = T)[1:10] # most common around 15

#registerDoMC(detectCores()-1)
mls_thin <- llply(1:n.indiv(ms_clean), .fun=function(i){
  print(paste0("Working on animal ",i," of ",n.indiv(ms_clean)))
  m <- ms_clean[[i]]
  id <- paste(i,m@idData$Species,m@idData$Tag.Type,m@idData$Animal.ID,m@idData$ID, sep="_")
  summary(timeLag(m))

  m_thin <- thinTrackTime(m, interval = as.difftime(15, units='mins'),
                          tolerance = as.difftime(5, units='mins'))
  # Add the thinned segment selection as a column to the dataset
  m_thin$thin_segm <- c(NA, as.character(m_thin@burstId))
  #summary(timeLag(m_thin,"mins")[m_thin@burstId=="selected"])
  if(n.locs(m_thin)>0){
    #summary(m_thin@burstId); table(m_thin$thin_segm)
    #m_thin is a burst, so associated to each segment there is "selected" or "not selected"
    # We first calculate speed, distances and angles, not to change the sequence of segments
    m_thin$tlag_m <- c(NA, move::timeLag(m_thin,"mins"))
    m_thin$moveSpeed <- c(NA, move::speed(m_thin))
    m_thin$moveDistance <- c(NA, move::distance(m_thin))
    m_thin$vertDist <- c(NA, m_thin$z[-1] - m_thin$z[-length(m_thin$z)])
    m_thin$climbRate <- m_thin$vertDist/m_thin$tlag_m
```

```
    #  m_thin$glideRatio <- m_thin$moveDistance/m_thin$vertDist
    m_thin$turnAngle <- c(NA, turnAngleGc(m_thin), NA)
    m_thin$heading <- c(NA, angle(m_thin))
    #summary(m_thin$tlag_m[m_thin$thin_segm=="selected"])
    #summary(m_thin$tlag_m[m_thin$thin_segm=="notSelected"])
    return(m_thin)}
}, .parallel=F)

## save the list of move burst objects with the selected/unselected segments and flight parameters
save(mls_thin, file="out2_ctmm_vulture_moveburst_noOutliers_wholTraj_thinNOthin_flightPar.rdata")


# Create a data.frame with only "reliable" speeds/distances ####
#_________________________________________________________________

df_thin_rel <- do.call(rbind, llply(1:length(mls_thin), .fun=function(i){
  print(paste0("Working on animal ",i," of ",length(mls_thin)))
  m <- mls_thin[[i]]
  id <- paste(i,m@idData$Species,m@idData$Tag.Type,m@idData$Animal.ID,m@idData$ID, sep="_")
  # When transforming into a dataframe the burstID gets automatically added to the dataframe
  df_thin <- as.data.frame(m)
  # But not @idData so we add it
  df_thin <- cbind(df_thin, m@idData)
  # Remove unnecessary/duplicated columns
  df_thin <- df_thin[,!names(df_thin)%in%c("coords.x1","coords.x2", "optional",
                                          "sensor","timestamps","burstId")]
  # And finally filter only the "selected" segments (segments with the 15+-5 min timelag)
  df_thin <- df_thin[which(df_thin$thin_segm=="selected"),]
  #table(df_thin$thin_segm)
  #summary(df_thin$tlag_m)
  # And keep only segments with low variance in speed
  if(nrow(df_thin[which(df_thin$VAR.speed <= 0.5),])>0){
    df_thin_sub <- df_thin[which(df_thin$VAR.speed <= 0.5),]
    print(summary(df_thin_sub$tlag_m))
    print(summary(df_thin_sub$outlieSpeed))
    print(summary(df_thin_sub$moveSpeed))
    return(df_thin_sub)
  }
}))
df_thin_rel_cv <- df_thin_rel[df_thin_rel$Species == "CV",]
# Save output file with thinned and reliable data for Cape Vulture for next analyses
save(df_thin_rel_cv, file="out2_ctmm_vulture_dataset_thin15min_flightPar_reliableSpeeds_onlyCV.rdata")
```

### Step 3 - Daily flight parameters

```
library(lubridate)

path <- "/home/mscacco/ownCloud/VulturesMPI-VulProKerry/"
setwd(path)

# Load the vulture thinned data for the species CV, with only "reliable speeds"
#(data thinned to 15+-5 min and VAR.Speed <= 0.5), object df_thin_rel_cv
load("out2_ctmm_vulture_dataset_thin15min_flightPar_reliableSpeeds_onlyCV.rdata")
```

```
# Create a list of two elements (one per tag type) - movestack of individuals per tag type
tagType_ls <- split(df_thin_rel_cv, df_thin_rel_cv$Tag.Type)
names(tagType_ls)


# Create a DF with one entry per date per ID ####
#_________________________________________________
# and associated daily information, such as
#cumulative daily distance and diameter, prop of time spent flying, days from deployment

traj_ls <- split(df_thin_rel_cv, df_thin_rel_cv$ID)

# Lapply to calculate daily parameters
allInd_dailyDF <- do.call(rbind, lapply(traj_ls, function(m){
  message(paste0("Working on trajectory ",unique(m$ID)))
  # Extract dates
  m$dates <- as.character(date(m$timestamp))
  # Subset only actual flight locations
  m_sub <- m[which(m$moveSpeed >= 2),]
  # daily movement extent (min and max lon lat)
  minCoords <- aggregate(m[,c("longitude","latitude")], by=list(m$dates), FUN="min")[,2:3]
  maxCoords <- aggregate(m[,c("longitude","latitude")], by=list(m$dates), FUN="max")[,2:3]
  # First calculate parameters for the entire dataset (m)
  df1 <- data.frame(Date=unique(m$dates),
                    unique(m[,c("ID","Animal.ID","Species","Logger.Type","Group","Tag.Type",
                                "Tag.ID","Sex","Age.Class","Deployment.Date","Comment")]),
                    NdaysFromDeployment=as.numeric(difftime(
                      as.Date(unique(m$dates), format="%Y-%m-%d"),
                      as.Date(unique(m$Deployment.Date), format="%d/%m/%Y"), units='days')),
                    Nloc=as.numeric(table(m$dates)),
                    cumDailyDiam_km=pointDistance(p1=minCoords, p2=maxCoords, lonlat=T)/1000,
                    sumReliableTimelags_h=(aggregate(m$tlag_m, by=list(m$dates),
                                                     FUN="sum", na.rm=T)[,2])/60,
                    start_timestamp=aggregate(m$timestamp, by=list(m$dates),
                                              FUN="min", na.rm=T)[,2],
                    end_timestamp=aggregate(m$timestamp, by=list(m$dates),
                                            FUN="max", na.rm=T)[,2],
                    trackingDuration_h=aggregate(m$timestamp, by=list(m$dates),
                                                 FUN=function(x){difftime(
                                                   max(x),min(x), units='hours')})[,2],
                    Ndays_tracking=length(unique(m$dates)))
  # Then calculate only information concerning the flight segments (m_sub)
  df2 <- data.frame(Date=unique(m_sub$dates),
                    Nloc_flight=as.numeric(table(m_sub$dates)),
                    cumDailyDist_km=(aggregate(m_sub$moveDistance, by=list(m_sub$dates),
                                               FUN="sum", na.rm=T)[,2])/1000,
                    med_flightAlt=aggregate(m_sub$z, by=list(m_sub$dates),
                                            FUN="median", na.rm=T)[,2],
                    max_flightAlt=aggregate(m_sub$z, by=list(m_sub$dates),
                                            FUN="max", na.rm=T)[,2],
                    med_flightSpeed=aggregate(m_sub$moveSpeed, by=list(m_sub$dates),
                                              FUN="median", na.rm=T)[,2],
                    sumFlightTimelags_h=(aggregate(m_sub$tlag_m, by=list(m_sub$dates),
```

```
                                                    FUN="sum", na.rm=T)[,2])/60)
  # Merge the two datasets and calculate some extra infos
  dailyDF <- merge(df1, df2, by="Date", all.x=T)
  dailyDF$sumFlightTimelags_h[which(is.na(dailyDF$sumFlightTimelags_h))] <- 0
  dailyDF$Nloc_flight[which(is.na(dailyDF$Nloc_flight))] <- 0
  dailyDF$prop_hFlightTime <- dailyDF$sumFlightTimelags_h/dailyDF$sumReliableTimelags_h
  dailyDF$Ndays_flight <- length(unique(m_sub$dates))
  dailyDF$anyFlight <- 0
  dailyDF$anyFlight[dailyDF$sumFlightTimelags_h>0] <- 1
  return(dailyDF)
}))
write.csv(allInd_dailyDF, "dailySummary_perDate_perTrackingID_onlyCV_thin.csv", row.names = F)
save(allInd_dailyDF, file="dailySummary_perDate_perTrackingID_onlyCV_thin.rdata")

# Create a DF with one entry per trajectory/individual, and associated average information ####
#_______________________________________________________________________________________________
# such as avg cumulative daily distance and diameter, median of the speed and altitude etc

if(dir.exists("DescriptiveSummaryStats_april2020_onlyCV_ctmm_thin")==F){
  dir.create("DescriptiveSummaryStats_april2020_onlyCV_ctmm_thin")}

setwd(paste0(path,"DescriptiveSummaryStats_april2020_onlyCV_ctmm_thin"))

traj_dailyls <- split(allInd_dailyDF, allInd_dailyDF$ID)

allInd_avgDF <- do.call(rbind, lapply(traj_dailyls, function(m){
  newDF <- data.frame(unique(m[,2:12]),
                      med_flightAlt=median(m$med_flightAlt, na.rm=T),
                      max_flightAlt=max(m$max_flightAlt, na.rm=T),
                      med_flightSpeed=median(m$med_flightSpeed, na.rm=T),
                      avg_cumDailyDist_km=mean(m$cumDailyDist_km, na.rm=T),
                      avg_cumDailyDiam_km=mean(m$cumDailyDiam_km, na.rm=T),
                      avg_prop_hFlightTime=mean(m$prop_hFlightTime, na.rm=T),
                      start_timestamp=min(m$start_timestamp),
                      end_timestamp=max(m$end_timestamp),
                      trackingDuration_d=as.numeric(difftime(
                        max(m$end_timestamp),min(m$start_timestamp), units='days')),
                      Nloc=sum(m$Nloc),
                      Nloc_flight=sum(m$Nloc_flight),
                      Ndays_tracking=unique(m$Ndays_tracking),
                      Ndays_flight=unique(m$Ndays_flight))
  return(newDF)
}))
write.csv(allInd_avgDF, "summaryInfo_perTrackingID_onlyCV_thin.csv", row.names = F)
```

### Step 4 - Statistical models

```
# Import data ####
#________________

path <- "/home/mscacco/"
setwd(paste0(path,"ownCloud/VulturesMPI-VulProKerry"))
```

```
library(mgcv) #for gams
library(lme4)
library(MASS)
library(data.table)
library(ggplot2)
source(paste0(path,"ownCloud/Martina/PHD/R_functions/standard_error.R"))

# Read in the df with the info summarized per date
df <- read.csv("dailySummary_perDate_perTrackingID_onlyCV_thin.csv", as.is=T)
df$Date <- as.Date(df$Date)
df$Tag.Type <- factor(df$Tag.Type, levels=c("Legband","Patagial"))
df$Group <- factor(df$Group, levels=c("WILD","CB"))
df$Animal.ID <- as.factor(df$Animal.ID)
df$Age.Class <- factor(df$Age.Class, levels=c("Fledging", "Juvenile", "Subadult", "Adult"))
df$Julian <- as.numeric(format(df$Date, "%j"))
df <- df[df$NdaysFromDeployment>=0,] # Remove days before deployment date (15 entries)

table(df$anyFlight)
table(df$anyFlight, df$Tag.Type)
table(df$anyFlight)/nrow(df)
mean(sapply(split(df$trackingDuration_h, df$ID), sum))
stderr(sapply(split(df$trackingDuration_h, df$ID), sum))
range(sapply(split(df$trackingDuration_h, df$ID), sum))

mean(df$prop_hFlightTime, na.rm=T)
stderr(df$prop_hFlightTime)

mean(df$med_flightSpeed, na.rm=T); stderr(df$med_flightSpeed)
mean(df$cumDailyDist_km, na.rm=T); stderr(df$cumDailyDist_km)

#time range
range(df$start_timestamp[df$Tag.Type=="Legband"])
range(df$start_timestamp[df$Tag.Type=="Patagial"])
table(df$Tag.Type)


# GAMMs ####
------------

# response: daily probability of flight ----
#____________________________________________
# days with movement vs days with no movement (0|1) (threshold 2 m/s)
df_cc <- df[complete.cases(df[,c("anyFlight","Tag.Type","NdaysFromDeployment
                                 Group","Animal.ID")]),]

# flight prob. is calculated on both flight and non flight locations so we use "all" locations
gamProb <- gamm(anyFlight ~ Tag.Type * Group +
                  s(NdaysFromDeployment) +
                  s(Nloc) +
                  s(Julian,bs="cc"),
                random = list(Animal.ID=~1),
                data=df_cc,
                method="REML",
```

```
                family=binomial(link="logit"))
summary(gamProb$gam)
summary(gamProb$lme)
plot.gam(gamProb$gam, page=1)
par(mfrow=c(2,2))
gam.check(gamProb$gam)
# residuals accounting for random terms
gamRes <- resid(gamProb$lme, type = "normalized")
acf(gamRes)
hist(gamRes)

## Extract probabilities of flight for better interpretation
# plogis(coef) is same as doing: 1 / (1 + exp(- coef))
modcoefs <- coefficients(gamProb$gam)
plogis(modcoefs["(Intercept)"]) #prob for wild individuals with legbands
plogis(sum(modcoefs[c("(Intercept)","Tag.TypePatagial","GroupCB",
                      "Tag.TypePatagial:GroupCB")]))#prob for captive individuals with patagial
## Extract random structure
#intercept for each individual
gamProb$lme$coefficients$random$Animal.ID
RIsd=1.42
plogis(modcoefs["(Intercept)"]+(-1.96*RIsd))#range of intercept among individuals
plogis(modcoefs["(Intercept)"]+(1.96*RIsd))

# response: daily amount of flight ----
#_________________________________________
# daily proportion of flight/non flight timelags (moving/non moving) excluding 0s (threshold 2 m/s)
df_cc <- df[complete.cases(df[,c("prop_hFlightTime","Tag.Type","NdaysFromDeployment",
                                 "Group","Animal.ID")]),]
df_cc <- df_cc[df_cc$anyFlight == 1,] #only consider days with some movement happening

#summary of individuals' daily proportion of flying time
summary(sapply(split(df_cc, df_cc$Animal.ID), function(id) mean(id$prop_hFlightTime)))

# prop_hFlightTime was calculated as sumFlightTimelags_h/sumReliableTimelags_h
# Recalculate using minutes to avoid decimals in the weights
sumReliableTimelags_min <- round(df_cc$sumReliableTimelags_h*60)
sumFlightTimelags_min <- round(df_cc$sumFlightTimelags_h*60)
prop_FlightTime_min <- sumFlightTimelags_min/sumReliableTimelags_min

# using binomial with proportion and weights
gamFlight <- gamm(prop_FlightTime_min ~ Tag.Type * Group +
                   s(NdaysFromDeployment) +
                   s(Nloc_flight) +
                   s(Julian,bs="cc"),
                  random = list(Animal.ID=~1),
                  weights=sumReliableTimelags_min,
                  data=df_cc,
                  method="REML",
                  family=binomial(link="logit"))
summary(gamFlight$gam)
par(mfrow=c(1,1))
plot.gam(gamFlight$gam, page=1)
plot(gamFlight$gam$fitted.values, prop_FlightTime_min)
```

```
summary(gamFlight$lme)
par(mfrow=c(2,2))
gam.check(gamFlight$gam)
# residuals accounting for random terms
gamRes <- resid(gamFlight$lme, type = "normalized")
acf(gamRes)
hist(gamRes)

## Extract daily proportion of flight for better interpretation
modcoefs <- coefficients(gamFlight$gam)
plogis(modcoefs["(Intercept)"]) # for wild individuals with legbands
plogis(sum(modcoefs[c("(Intercept)","GroupCB")]))# for captive individuals with legband
plogis(sum(modcoefs[c("(Intercept)","Tag.TypePatagial")]))# for wild individuals with patagial
plogis(sum(modcoefs[c("(Intercept)","Tag.TypePatagial","GroupCB",
                      "Tag.TypePatagial:GroupCB")]))# for captive individuals with patagial
## Extract random structure
#intercept for each individual
gamFlight$lme$coefficients$random$Animal.ID
RIsd=0.26
plogis(modcoefs["(Intercept)"]+(-1.96*RIsd))
plogis(modcoefs["(Intercept)"]+(1.96*RIsd))

# response: daily cumulative distance (step lengths) in meters ----
#__________________________________________________________________
# Cumulative distance was calculated including only flight locations (with speed >= 2 m/s)
df_cc <- df[complete.cases(df[,c("cumDailyDist_km","Tag.Type","Nloc_flight",
                                 "NdaysFromDeployment","Group","Animal.ID")]),]

#summary of individuals' daily cumulative distance travelled
summary(sapply(split(df_cc, df_cc$Animal.ID), function(id) mean(id$cumDailyDist_km)))

gamDist <- gamm(sqrt(cumDailyDist_km) ~ Tag.Type * Group +
                   s(NdaysFromDeployment) +
                   s(Nloc_flight) +
                   s(Julian,bs="cc"),
                 random = list(Animal.ID=~1),
                 family = gaussian,
                 method="REML",
                 data=df_cc)
summary(gamDist$gam, cor = FALSE)
anova(gamDist$gam)
par(mfrow=c(1,1))
plot(gamDist$gam, page=1)
plot(gamDist$gam$fitted.values, sqrt(df_cc$cumDailyDist))
summary(gamDist$lme)
par(mfrow=c(2,2))
gam.check(gamDist$gam)
# residuals accounting for random terms
gamRes <- resid(gamDist$lme, type = "normalized")
acf(gamRes)
hist(gamRes)

## Extract values in Km for better interpretation (distance was sqrt so now we use ^2)
modcoefs <- coefficients(gamDist$gam)
```

```
(modcoefs["(Intercept)"])^2 # for wild individuals with legbands
(sum(modcoefs[c("(Intercept)","GroupCB")]))^2 # for captive individuals with legband
(sum(modcoefs[c("(Intercept)","Tag.TypePatagial")]))^2 # for wild individuals with patagial
(sum(modcoefs[c("(Intercept)","Tag.TypePatagial","GroupCB",
                "Tag.TypePatagial:GroupCB")]))^2 # for captive individuals with patagial
## Extract random structure
gamDist$lme$coefficients$random$Animal.ID
RIsd=0.37
(modcoefs["(Intercept)"]+(1.96*RIsd))^2
(modcoefs["(Intercept)"]+(-1.96*RIsd))^2


# response: daily median flight speed ----
#__________________________________________
# Median speed was calculated including only flight locations (with speed >= 2 m/s)
df_cc <- df[complete.cases(df[,c("med_flightSpeed","Tag.Type","Nloc_flight",
                                 "NdaysFromDeployment","Group","Animal.ID")]),]

gamSpeed <- gamm(sqrt(med_flightSpeed) ~ Tag.Type * Group +
                   s(NdaysFromDeployment) +
                   s(Nloc_flight) +
                   s(Julian,bs="cc"),
                 random = list(Animal.ID=~1),
                 family = gaussian, #Gamma(link="log"),
                 method="REML",
                 data=df_cc)
summary(gamSpeed$gam)
anova(gamSpeed$gam)
par(mfrow=c(1,1))
plot.gam(gamSpeed$gam, page=1)
plot(gamSpeed$gam$fitted.values, sqrt(df_cc$med_flightSpeed))
summary(gamSpeed$lme)
par(mfrow=c(2,2))
gam.check(gamSpeed$gam)
# residuals accounting for random terms
gamRes <- resid(gamSpeed$lme, type = "normalized")
acf(gamRes)
hist(gamRes)

## Extract values in m/s for better interpretation (speed was sqrt so now we use ^2)
modcoefs <- coefficients(gamSpeed$gam)
(modcoefs["(Intercept)"])^2 # for wild individuals with legbands
(sum(modcoefs[c("(Intercept)","GroupCB")]))^2 # for captive individuals with legband
(sum(modcoefs[c("(Intercept)","Tag.TypePatagial","GroupCB",
                "Tag.TypePatagial:GroupCB")]))^2 # for captive individuals with patagial
## Extract random structure
gamSpeed$lme$coefficients$random$Animal.ID
RIsd=0.10
(modcoefs["(Intercept)"]+(1.96*RIsd))^2
(modcoefs["(Intercept)"]+(-1.96*RIsd))^2
```
